# Circadian Control of Histone Turnover During Cardiac Development and Growth

**DOI:** 10.1101/2023.11.14.567086

**Authors:** Adrian Arrieta, Douglas J. Chapski, Anna Reese, Todd Kimball, Kunhua Song, Manuel Rosa-Garrido, Thomas M. Vondriska

**Author notes:** Correspondence: Thomas M. Vondriska Department of Anesthesiology & Perioperative Medicine David Geffen School of Medicine at UCLA CHS 37-100 650 Charles Young Dr. Los Angeles, CA 90095.

## Abstract

**Rationale:** During postnatal cardiac hypertrophy, cardiomyocytes undergo mitotic exit, relying on DNA replication-independent mechanisms of histone turnover to maintain chromatin organization and gene transcription. In other tissues, circadian oscillations in nucleosome occupancy influence clock-controlled gene expression, suggesting an unrecognized role for the circadian clock in temporal control of histone turnover and coordinate cardiomyocyte gene expression. **Objective:** To elucidate roles for the master circadian transcription factor, Bmal1, in histone turnover, chromatin organization, and myocyte-specific gene expression and cell growth in the neonatal period. **Methods and Results:** Bmal1 knockdown in neonatal rat ventricular myocytes (NRVM) decreased myocyte size, total cellular protein, and transcription of the fetal hypertrophic gene Nppb following treatment with increasing serum concentrations or the α-adrenergic agonist phenylephrine (PE). Bmal1 knockdown decreased expression of clock-controlled genes Per2 and Tcap, and salt-inducible kinase 1 (Sik1) which was identified via gene ontology analysis of Bmal1 targets upregulated in adult versus embryonic hearts. Epigenomic analyses revealed co-localized chromatin accessibility and Bmal1 localization in the Sik1 promoter. Bmal1 knockdown impaired Per2 and Sik1 promoter accessibility as measured by MNase-qPCR and impaired histone turnover indicated by metabolic labeling of acid-soluble chromatin fractions and immunoblots of total and chromatin-associated core histones. Sik1 knockdown basally increased myocyte size, while simultaneously impairing and driving Nppb and Per2 transcription, respectively. **Conclusions:** Bmal1 is required for neonatal myocyte growth, replication-independent histone turnover, and chromatin organization at the Sik1 promoter. Sik1 represents a novel clock-controlled gene that coordinates myocyte growth with hypertrophic and clock-controlled gene transcription.

## Introduction

During neonatal cardiac development in mammals, cardiomyocytes integrate myriad environmental cues such as changes in blood pressure, blood oxygen concentration, and circulating hormones with genetic programming to execute cardiac maturation (1). Part of this process includes a switch from cardiomyocyte proliferation to hypertrophy, which is accompanied by a loss of organ-level regenerative capacity within the first week after birth (2). Gene expression has been extensively studied during this period of cardiac maturation, as has the role of transcription factors and epigenetic modifiers in specifying myocyte cell fate. What remains unexplored is the role of chromatin organization during cardiac maturation. In this study, we sought to understand the role of histone turn over in myocyte growth during this critical neonatal period.

Since myocytes exit the cell cycle, they must switch from DNA replication-dependent to replication independent replacement of nucleosomes, the basic packaging units of chromatin. Much evidence suggests that histone replacement is a highly tuned phenomenon outside of the context of replication and that the process of histone replacement is coupled with transcription (3–5). In the adult heart, transcriptional responses to pathologic growth of pressure overload become entrained in chromatin, such that locus specific remodeling precedes later transcriptional adaptation (6). Also in the adult, nucleosome turnover (as measured by GFP-tagged histone H2B) is higher at actively transcribed genes, including those involved in cell identity and cardiac contractility, but also at many that are not cell type specific (3). This study also observed increased turnover at areas of active chromatin as indicated by histone marks such as H3 lysine 27 acetylation. However, many questions remained unanswered, including: what role does temporally precise histone and nucleosome turnover play earlier in development during the period right after myocyte proliferation wanes? To address this question, we examined core circadian clock proteins involved in cardiac development that potentially act through histones.

Circadian clocks function throughout the eukaryotic world to control rhythmic behaviors of multicellular organisms (7), and this rhythm can be influenced by external time cues, otherwise known as “zeitgebers”. The central clock housed in the suprachiasmatic nucleus in the brain is entrained by light (8), while the peripheral clock in the heart responds to feeding-fasting cycles and catecholamines (9–11). The molecular clock operates at the cell and organ level, underpinned by a subcellular network of transcriptional and protein circuits originally identified at the molecular level in flies with the discovery of the gene *Period* (12). Clock and Bmal1 are basic helix-loop-helix transcription factors that bind to E-box regulatory elements to promote transcription of a set of “clock-controlled” genes, including Cryptochrome (Cry1/Cry2) and Period (Per1/Per2/Per3). Cry and Per form heteromultimers with Casein kinase (CK) in the cytosol, inducing nuclear translocation, where the complex binds to Clock-Bmal1 heterodimers to repress their own transcription in a negative feedback loop, molecular features characteristic of circadian regulation across cells (13). In addition to transcriptional cycling, many intermediate steps controlling protein abundance have been shown to oscillate, including splicing, mRNA degradation, translation and protein degradation (14,15). Circadian rhythms participate in normal cardiac development and disease in vivo: altered circadian biology is associated with cardiac arrhythmias, myocardial infarction, hypertension and diabetes (16,17), and loss of Bmal1 in myocytes from birth causes dilated cardiomyopathy and metabolic derangement (18), although subcellular mechanisms are unclear. In non-cardiac tissues, Bmal1 has been implicated in daily oscillations of nucleosome occupancy, histone H3 acetylation, and RNA polymerase II (RNAPII) recruitment, coordinate with gene expression (19–21). In the present study, we sought to examine the role of Bmal1 in cell growth and chromatin remodeling during the neonatal period. We find that Bmal1 is critical for normal myocyte growth and histone turnover, linking circadian control of chromatin remodeling to normal cardiac maturation after birth.

## Results

### Depletion of Bmal1 loss myocyte hypertrophy and gene expression

To determine whether Bmal1 is required for myocyte hypertrophy in the absence of hemodynamic influence (22–24), cultured neonatal rat ventricular myocytes (NRVM) were transfected with siRNA targeting Bmal1 and then incubated with 0, 2, or 10% fetal calf serum (FCS) for 48Hr to dose-dependently assess serum-mediated myocyte growth (Fig. 1A). Cultures treated with 0% FCS were supplemented with ITS to maintain cell viability without stimulating hypertrophy (22,23). Incubation of cultured with 2% FCS significantly affected Bmal1 protein and transcript levels of Bmal1, but greater increases in FCS reversed this effect, indicating circadian regulation of clock gene expression at both transcript and protein levels in myocytes can be influenced by the presence of factors circulating in blood (Fig. 1B, 1C).

**Figure 1:**
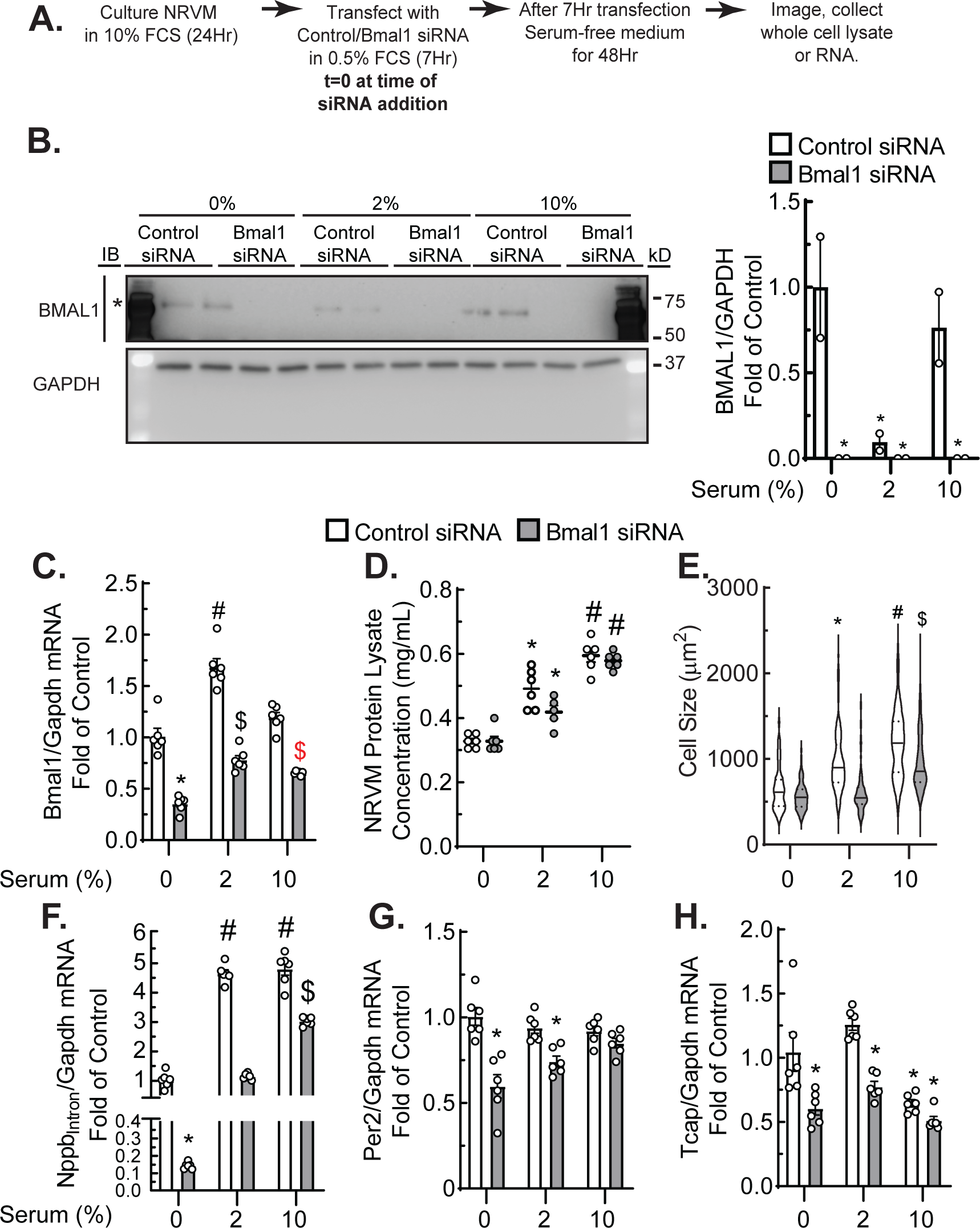
Bmal1 loss of function impairs expression of clock-controlled and hypertrophic genes. **A**) Experimental timeline of cultured NRVM transfected with scrambled or Bmal1-targeted siRNA and treated with increasing serum concentrations. **B, C**) Bmal1 and Gapdh immunoblots and RT-qPCR demonstrating Bmal1 knockdown. **D, E**) NRVM lysate protein concentration and cell size measurements. **F**-**H**) NppbIntron, Per2, and Tcap RT-qPCR. *, ^#^, ^$^ indicate difference from all other groups in the same treatment condition, p≤0.05 by two-way ANOVA followed by Tukey’s post hoc analysis.

Protein concentration of whole cell lysates increased as a function of serum concentration and was not significantly altered by Bmal1 knockdown (Fig. 1D). Under serum-free conditions Bmal1 knockdown did not significantly affect myocyte size: however, Bmal1 depletion was effective in preventing hypertrophy induced by low and high serum concentrations (Fig. 1E). These observations indicate that Bmal1 expression is dynamic and required at precise levels to engender a growth response: higher levels of serum (10 vs. 2%) did not induce more Bmal1 but did induce greater cell growth, and Bmal1 was necessary at both serum concentrations for this growth (Fig. 1C & E).

We next assessed the effects of Bmal1 depletion on expression of hypertrophic genes and on other clock-controlled genes in the setting of neonatal growth. Natriuretic peptide B (*Nppb*) is upregulated in hypertrophic myocytes in vivo and in culture: we specifically measured the levels of Nppb transcripts containing an intron (Nppbintron) to more accurately reflect *de novo* transcription of this gene in the early stages of myocyte hypertrophy (14,25). Under serum-free conditions, Bmal1 knockdown resulted in a 90% decrease in Nppb expression (Fig. 1F). Treatment with 2% or 10% serum induced robust increase in Nppb transcript levels, an effect partially attenuated by depletion of Bmal1 (Fig. 1F). With Bmal1 present, 2% FCS also resulted in a 4-fold increase in Nppb expression in control transfected cells. Whereas control transfected cells displayed no further increase in NppbIntron between 2% and 10% FCS, NRVM lacking Bmal1 showed a continued upregulation of NppbIntron. To examine other clock genes, we focused on Per2, as its regulation by Bmal1 has been interrogated in the heart and in other tissues (19,20), and on titin-cap (TCAP), which is an established striated muscle-specific clock-controlled gene (26). Bmal1 knockdown significantly decreased abundance of Per2 under basal conditions or following 2% FCS, with no change observed in Per2 expression at 10% FCS (Fig. 1G). A similar pattern of Tcap expression was observed: Bmal1 knockdown significantly decreased Tcap expression at 0 and 2% FCS, with no significant difference in Tcap expression change noted with 10% FCS, though 10% FCS also decreased Tcap expression compared to 0 and 2% FCS (Fig. 1H). These results suggest that Bmal1 is required for myocyte hypertrophy and the circadian clock regulates the level of hypertrophic gene transcription in neonatal myocytes.

### Replication-independent histone H3.3a expression is critical for myocyte growth and gene expression

Neonatal myocytes undergo limited cell division turnover in culture (24)—thus, we hypothesized that the majority of histone turnover occurs in a replication-independent manner, likely concomitant with *de novo* gene transcription (4). To determine whether replication-independent histone variants are required for myocyte-specific fetal hypertrophic gene expression, NRVM were transfected with an siRNA targeting the replication independent histone variant H3.3a, which is exchanged on chromatin during transcription (4,5). This approach produced a protein level decrease of 50% for H3.3 and 25% for total H3 (Figs. 2A, 2B; RT-qPCR shows this depletion is specific for H3.3a, Fig. 2C). Knockdown of H3.3a impaired hypertrophic gene transcription and myocyte hypertrophy, as indicated by decreased NppbIntron and myocyte size (Fig. 2D), consistent with the prior notion (4) that replication-independent histone H3.3 turnover is critical for *de novo* fetal hypertrophic gene transcription in myocytes, as well as for myocyte hypertrophy.

**Figure 2:**
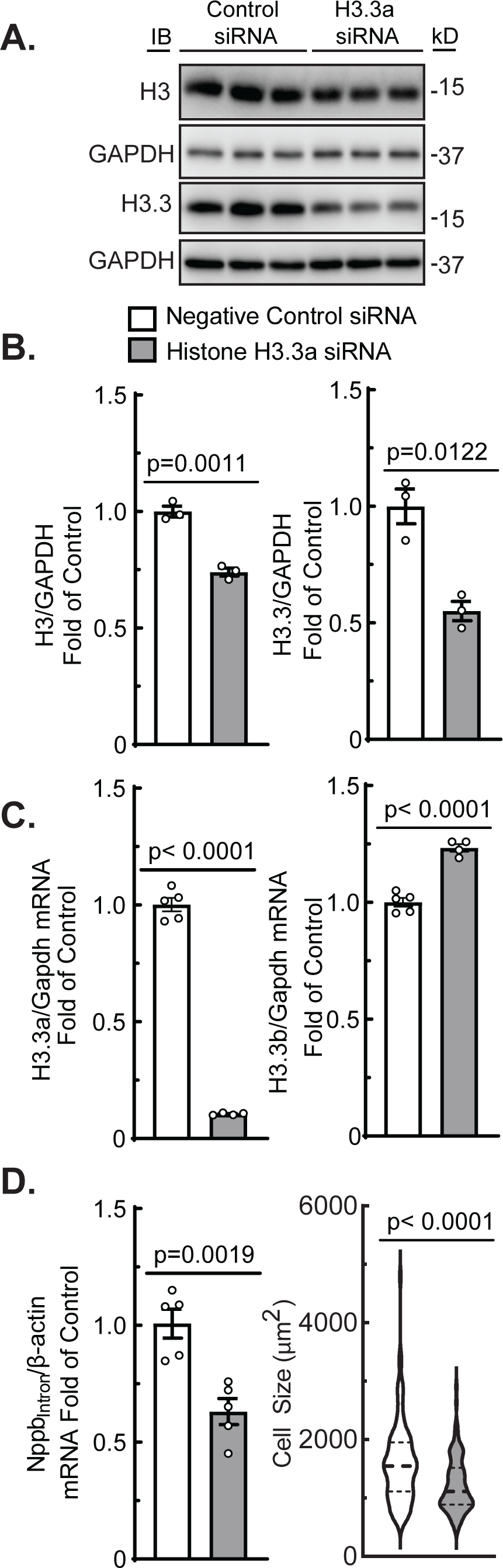
Replication-independent histone turnover is critical for myocyte growth and hypertrophic gene expression. Cultured NRVM were transfected with siRNA against H3.3a and knockdown was confirmed by immunoblot (**A-B**). H3.3a knockdown results in decreased H3.3a but not H3.3b transcript levels (**C**), and a decrease in NppbIntron and cell size (**D**). P-values indicated are generated by unpaired t-tests.

### Bmal1 loss of function disrupts histone stoichiometry and impairs histone turnover

Having linked histone abundance with myocyte growth, we next sought to definitively test whether Bmal1 regulates histone expression and assembly into chromatin. In this experiment, cultures were incubated in serum-free medium for 72 hours to minimize the effects of serum-borne factors that can affect clock activity (Fig. 3A). To determine whether Bmal1 knockdown impairs hypertrophy via disruption of histone turnover, we assessed the relative levels of histone H2A, H2B, H3 and H4, in whole cell lysate and in acid soluble chromatin fractions relative to GAPDH and total protein (Oriole staining), respectively. Depletion of Bmal1 led to a decrease in both total protein levels and cell size (Fig. 3B). When controlling for total protein abundance, the levels of H3 and H2A, but that of neither H2B nor H4, were increased following Bmal1 knockdown (Fig.3C). We also noted the presence of a lower molecular weight band in our H3 and H2B immunoblots, with significant accumulation of H3 after Bmal1 knockdown (H314kD, H2B14kD; Fig. 3D).

**Figure 3:**
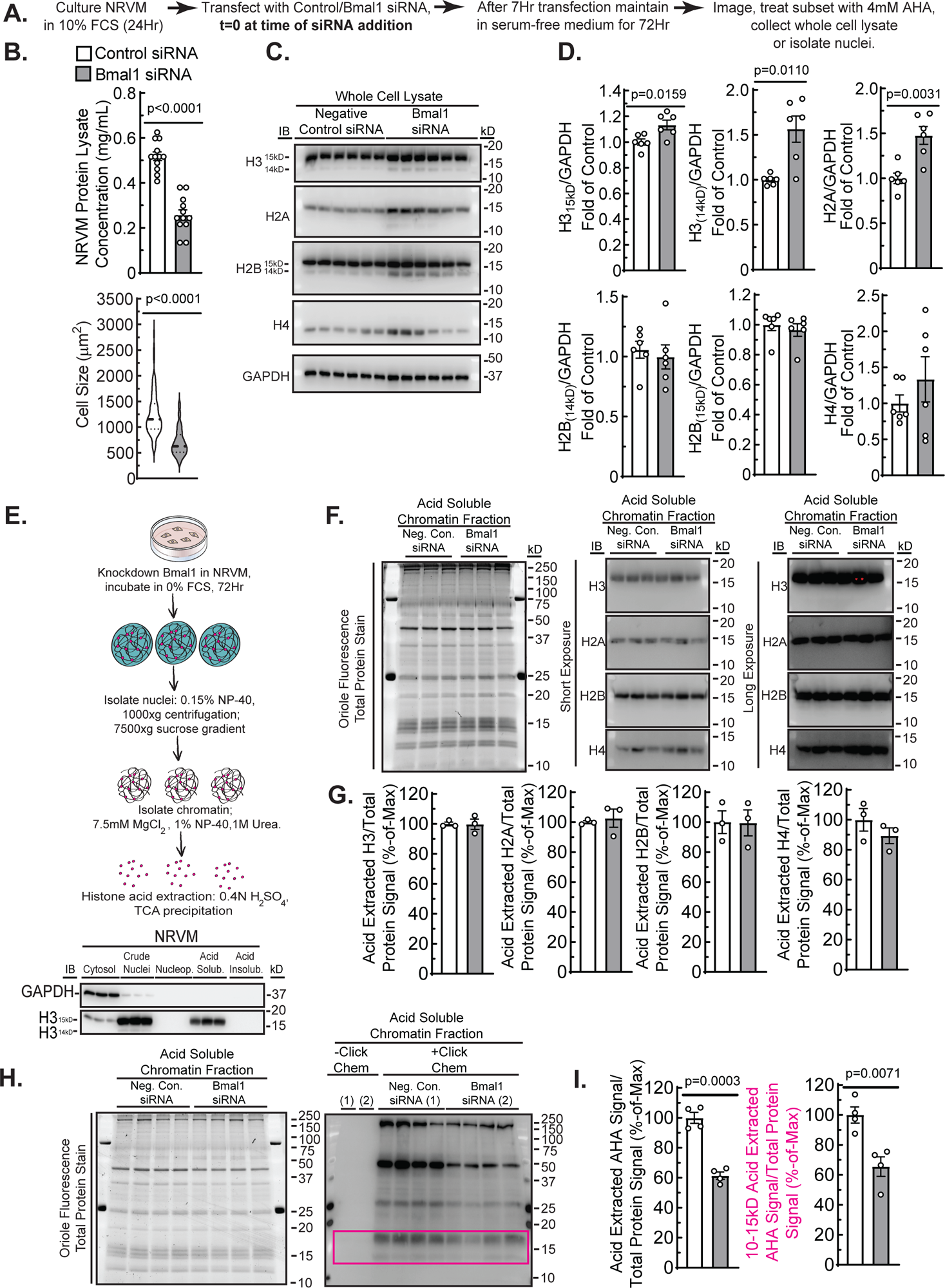
Bmal1 loss of function disrupts histone stoichiometry and impairs histone turnover. **A)** Experimental timeline of cultured NRVM transfected with scrambled or Bmal1-targeted siRNA, maintained in serum-free medium for 72Hr and, where indicated, treated with 4mM AHA for 4Hr. **B**) NRVM protein lysate concentration and cell size measurement of control or Bmal1 siRNA-transfected cells. **C**) NRVM protein lysates were probed for core histone levels and GAPDH via immunoblot and quantified in **D**). Indicated p-values were generated by student’s unpaired t-tests. **E**) Diagram of nuclei isolation from NRVM, followed by isolation and acid extraction of chromatin. Isolation of chromatin-associated histones was validated by sub-nuclear fractionation and H3 and GAPDH immunoblots. **F**) Oriole fluorescence protein stain and H3, H4, H2A, H2B immunoblots of acid-soluble chromatin fractions. Immunoblot signal was normalized to total protein oriole fluorescence signal from NRVM transfected with control or Bmal1-targeted siRNA, with quantifications shown in **G**). **H**) Oriole fluorescence staining and streptavidin-HRP blot of acid-soluble chromatin fractions following click-chemistry with biotin-alkyne. **I**) Quantification of total AHA-labeled acid soluble chromatin fractions, normalized to oriole fluorescence. Indicated p-values were generated via student’s unpaired t-tests.

We next sought to distinguish total histone levels from those associated with chromatin in the nucleus, using an established subnuclear fractionation method (27). This approach revealed H314kD to be associated with intact nuclei, but not with acid soluble chromatin fractions, suggesting this band reflects H3 that has not been incorporated into chromatin (Figs. 3C-E). Furthermore, we found no differences in histone abundance at the level of chromatin (Fig. 3F&G), suggesting that differences in total abundance of cellular histones are maintained in pools of histones not bound to chromatin, i.e., as part of the pool available for nucleosome exchange.

To determine if, at the chromatin level, there is decreased histone turnover 72 hours post-transfection, NRVM were metabolically labeled with L-azidohomoalanine (AHA) for 4Hr, after which acid soluble chromatin fractions were isolated and subjected to click chemistry with biotin-alkyne followed by SDS-PAGE and streptavidin-HRP blotting (28). Consistent with the observed decrease in total cellular protein following Bmal1 knockdown, streptavidin-HRP blots revealed significantly less metabolic labeling of acid soluble chromatin fractions with Bmal1 knockdown (Fig. 3H&I). Additionally, Bmal1 depletion significantly decreased the levels of low molecular weight bands indicated in (Fig. 3H), where AHA-labeled histones have previously been shown to migrate(28).

To support the notion that these low molecular weight bands are histones and the observed effect of decreased abundance is indicative of impaired histone turnover on chromatin following Bmal1 depletion, neonatal rat ventricular fibroblasts were subjected to 72 hours of serum starvation followed by treatment with 20% FCS to stimulate synchronous cell cycle progression, i.e., replication-dependent histone turnover (29). Cultures were then labeled with 4mM AHA at 5Hr intervals followed by click chemistry with biotin-alkyne on acid soluble chromatin fractions (Supplemental Fig. 1A&B), where we observed accumulation of low molecular weight bands migrating at 15kD, 10-15Hr post-treatment with 20% FCS (Fig. S1C). Next, we performed pull downs with streptavidin-conjugated beads, followed by immunoblotting for histone H3. Finally, parallel cultures treated with serum for 15Hr revealed a significant increase in cyclin A2, which is required for entry into and progression of S-phase (29,30), and therefore replication-dependent histone turnover (Fig. S1D-E). The observations that Bmal1 knockdown results in 1) accumulation of un-chromatinized histones, 2) no change in the levels of chromatin-associated histones, and 3) decreased metabolic labeling of histones, in conjunction with results presented in Fig. S1, demonstrate that we are detecting labeled and therefore newly synthesized/chromatinized histones, support the notions that we are measuring active histone turnover at the chromatin level, and that Bmal1 is a critical mediator of histone turnover.

### Salt-inducible kinase 1 is a novel clock-controlled gene induced in response to growth stimuli

Having shown that Bmal1 acts to remodel chromatin and influence transcription in the post-natal period, we next sought to identify myocyte-specific Bmal1 target gene(s) that can influence both myocyte growth and entrainment of the myocyte clock in response to growth stimuli. We examined a transcriptomic study of embryonic and adult mouse myocytes (31) and compared genes significantly upregulated in the adult heart with experimentally determined regions of Bmal1 binding as measured by ChIP-seq (19). When analyzed by GProfiler, the genes occupied by Bmal1 across organs tended to be enriched in pathways associated with circadian regulation (Fig. 4A&B), including rhythmic biological processes, clock entrainment, and photoperiodism. Furthermore, those Bmal1 peaks specific to the heart were enriched for processes related to muscle cell differentiation and development and sarcomere assembly (Fig. 4C). We opted to focus on the gene encoding salt inducible kinase (Sik1), which has been previously shown to mediate myocyte hypertrophy in response to a growth stimulus in the adult heart (32) and found to contribute to circadian entrainment of the suprachiasmatic nucleus in response to light or serum shock (8), satisfying the criteria of a clock-controlled gene that can influence myocyte growth and entrainment of the myocyte clock. From our initial gene ontology analysis, it was not immediately clear how Sik1 could paradoxically be regulated by Bmal1 across organs and also be a cardiac-specific clock-controlled gene. To explore this gene further, we examined the Bmal1 ChIP-seq data (6,19) generated from heart, kidney, and liver with previously published ATAC-seq data from mouse hearts subjected to transverse aortic constriction-induced cardiac hypertrophy (6,19). This analysis revealed two Bmal1 binding sites, which co-localize with a cardiac-specific pattern of chromatin accessibility at the Sik1 promoter (Fig. 4D). RNA-seq data from the same mouse model of pressure overload shows decreased levels of Sik1 chromatin accessibility and transcripts as a function of time following injury (Fig. 4D), consistent with previous reports demonstrating that pressure overload can suppress clock-controlled gene expression (10). Additionally, in a separate dataset containing Mef2A, Nkx2.5, Tbx5, Srf, Gata4, H3K27Ac and RNAPII ChIP-seq data from embryonic and adult mouse hearts (31), we observed greater Sik1 promoter binding of RNAPII and Gata4 in the adult versus embryo (Fig. 4E). Because it has been shown that Gata4 transcriptional activity increases following a growth stimulus i.e. phenylephrine (PE) (33), we examined a published ATAC-Seq dataset (34) examining the effect of PE on NRVM chromatin— the same model system used in this study. This study observed two ATAC-seq peaks upstream of the Sik1 promoter (Fig. 4F), mimicking the finding from the adult mouse heart (Fig. 4D), and confirming cell type-specific, growth stimulus-induced chromatin remodeling at this locus in two separate species (rat and mouse).

**Figure 4:**
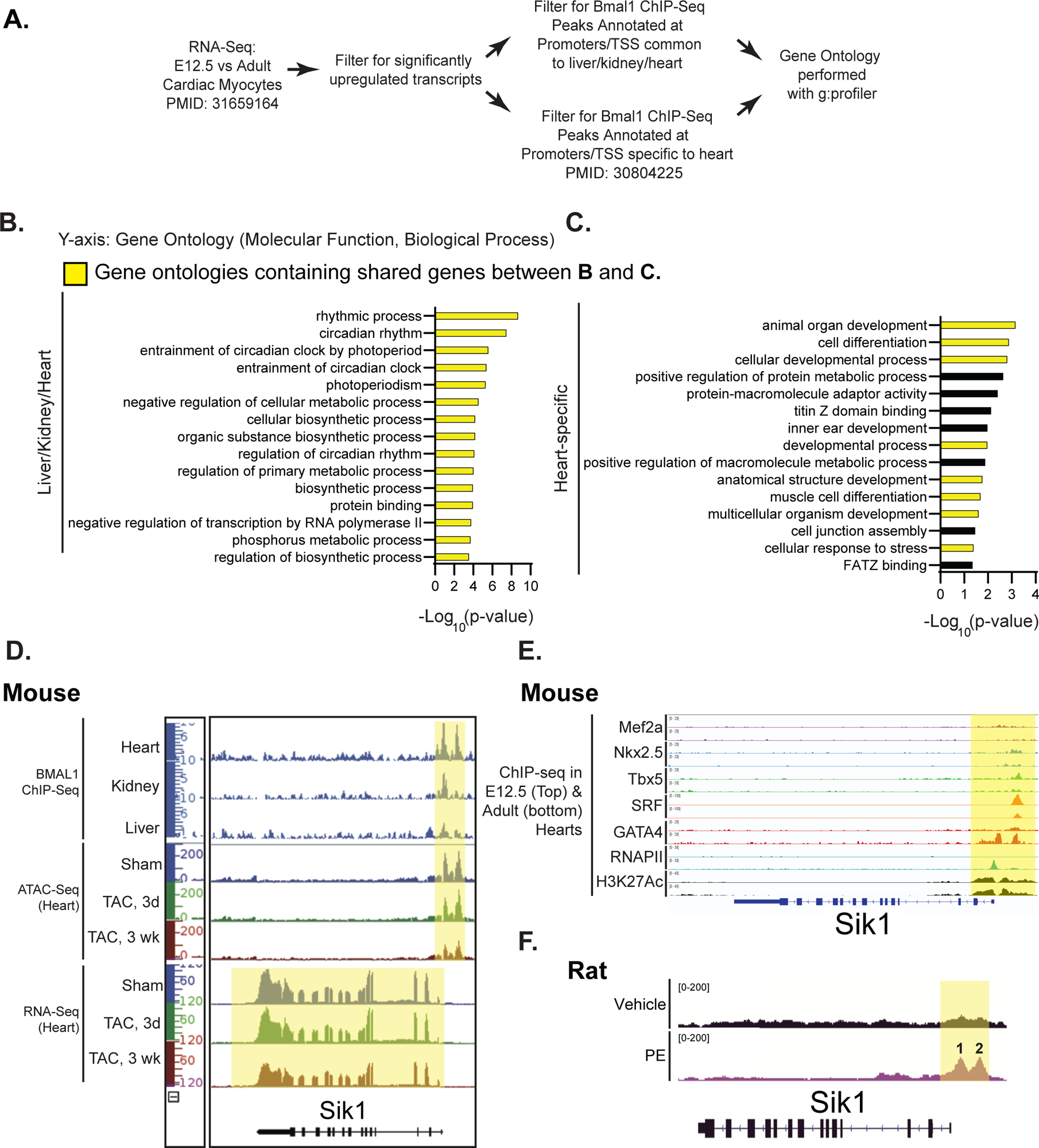
Salt-inducible Kinase 1 is a novel clock-controlled gene that is induced in response to a growth stimulus. **A**) Diagram of transcriptomic analysis to identify Bmal1 targets that are upregulated in the adult vs. embryonic heart, and to identify cardiac-specific Bmal1 target genes. **B**, **C**) G: profiler analysis indicating **B**) Bmal1-regulated gene ontologies common to the liver, kidney, heart, and **C**) Bmal1-regulated gene ontologies specific to the heart. **D**) Overlap of BMAL1 ChIP-Seq data (Beytebiere *et al.*, *Genes Dev*, 2019) from indicated mouse tissues and ATAC-seq and RNA-seq data of Sik1 from hearts of mice subjected to transaortic constriction (TAC)-mediated cardiac hypertrophy (Chapski *et al.*, *JMCC*, 2021). **E**) Overlay of Mef2A, Nkx2.5, Tbx5, SRF, GATA4, RNAPII, and H3K27Ac ChIP-Seq data of Sik1 from embryonic and adult mouse hearts (Akerberg *et al*., Nat Commun. 2019). **F**) ATAC-seq data of the Sik1 loci generated from NRVM, demonstrating increased Sik1 chromatin accessibility after treatment with PE (Golan-Lagziel *et al.*, *JMCC*, 2019).

### Bmal1 is required for Sik1 promoter accessibility and induction in response to a growth stimulus

We next sought to determine the chromatin remodeling mechanisms underpinning Bmal1-dependent regulation of Sik1 in the setting of cardiac maturation. Having shown that Bmal1 regulates Sik1 expression and that chromatin is remodeled at the Sik1 promoter in response to a growth stimulus, we employed a promoter accessibility assay to directly measure chromatin compaction (35). An MNase dose response was performed, whereby digestion with 0.1U MNase yielded a DNA laddering pattern indicative of partial digestion that was similar between cultures transfected with scramble or Bmal1-targeted siRNA (Fig. 5B). This result suggests that global chromatin accessibility is not affected by Bmal1 loss of function.

**Figure 5:**
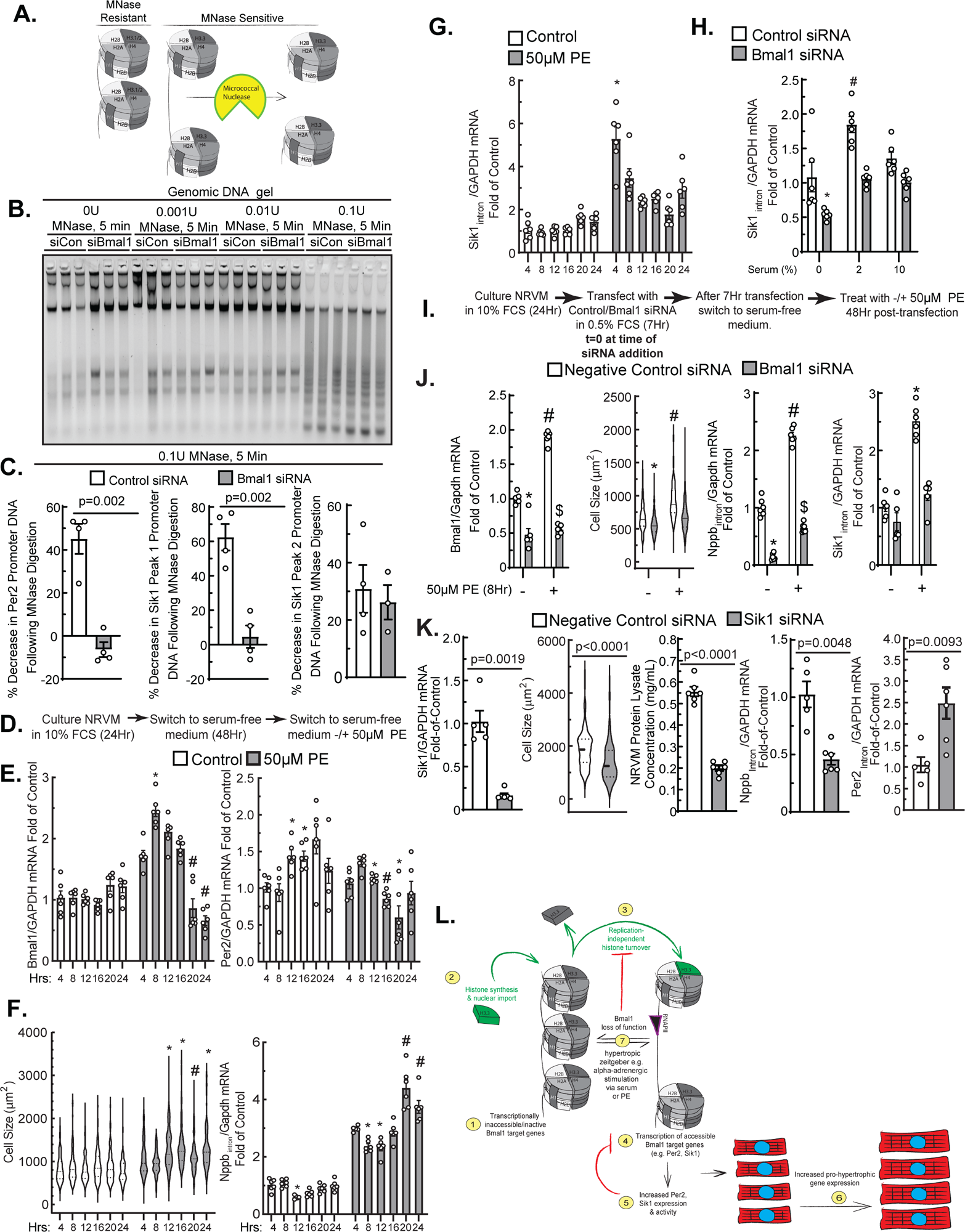
Bmal1 is required for Sik1 promoter accessibility and induction in response to a growth stimulus. **A)** Diagram of how MNase cleaves open regions of chromatin (MNase sensitive) as opposed to more compact regions (MNase resistant). **B**) Optimization of MNase for examination of chromatin compaction. 0.1U for 5min led to uniform digestions; this global pattern was unaffected by Bmal1 depletion. **C**) PCR after MNase reveals targeted decrease in chromatin accessibility after Bmal1 depletion in NRVM at Per2 and one of two Sik1 loci highlighted in Fig. 5E. Indicated p-values are generated by unpaired t-tests. **D**) Experimental timeline of cultured NRVM treated with -/+ 50μM phenylephrine (PE) for 4, 8, 12, 16, 20, or 24Hr. **E**) Clock-controlled gene transcripts measured by RT-qPCR included Bmal1 and Per2. **F**) Hypertrophy was assessed by measurement of cell size and nascent Nppb transcript levels (NppbIntron). *, # indicates difference from all other groups in the same treatment condition, p≤0.05 by one-way ANOVA followed by Tukey’s post hoc analysis. **G**, **H**) De novo Sik1 transcription (Sik1intron) was assessed by qPCR as in **E**) following treatment with PE (**G**) or increasing concentrations of serum as in Fig. 2F**-H** (**H**). For **E**-**G**, * indicates difference from all other groups in the same treatment condition, p≤0.05 by one-way ANOVA followed by Tukey’s post hoc analysis. For **H**, *, # indicates differences from all other groups by two-way ANOVA followed by Tukey’s post-hoc analysis. **I**) Experimental timeline of cultured NRVM treated with 50μM PE for 8Hr, 48Hr post-transfection with control siRNA or siRNA targeting Bmal1 (**J**). (**K**) 96Hr after Sik1 knockdown, myocyte hypertrophy was assessed by Nppbintron RT-qPCR and myocyte cell size and total cellular protein measurements, along with Per2intron qPCR to examine clock-controlled gene transcription. Indicated p-values were generated via student’s unpaired t-test. **L**) Summary diagram: In response to a hypertrophic stimulus, Bmal1 binds to transcriptionally inactive and inaccessible target genes (1) and activates their transcription via coordination of histone turnover (2–5). Sik1 then executes its pro-hypertrophic function, while Per2 serves to negatively regulate Bmal1 transcriptional activity (5–6), and Bmal1 loss-of-function maneuvers impair this signaling cascade (7).

As a positive control, we tested Per2, known to be regulated in a circadian manner dependent on Bmal1. MNase-qPCR of the DNA isolated from nuclei digested with 0.1U MNase revealed that control transfected NRVM displayed approximately 40% less Per2 promoter DNA levels following MNase digestion, whereas Bmal1 knockdown resulted in essentially no change in Per2 promoter DNA levels following digestion (Fig. 5C). This indicates that the normal role of Bmal1 is to keep this locus open. We next examined the two peaks of accessibility identified from prior examination of the Sik1 locus, which we have deemed peak 1 and peak 2 (Fig. 4F). We observed a 60% decrease in Sik1 promoter peak 1 DNA in control-transfected cells following MNase-qPCR and, similar to the Per2 promoter, depletion of Bmal1 abolished the actions of MNase to recover this locus, suggesting decreased accessibility of the Sik1 promoter peak 1. In contrast, peak 2 in the Sik1 promoter was impervious to the presence of Bmal1 (Fig. 5C). This may perhaps be due to differences in DNA base composition of target DNA, which can influence digestion by MNase (36), or differences in other chromatin binding factors occupying this site. It is intriguing to speculate that peak 1, given that it is not cardiac specific (Fig. 4D), may be of less importance to cell type specific regulation, but targeted disruption of the locus would be needed to conclusively test this possibility.

It has been previously shown that treatment of isolated cardiac myocytes with the catecholamine norepinephrine drives both myocyte hypertrophy and oscillatory expression of clock-controlled genes (11,23). Since expression of clock-controlled genes increases between the embryonic and the adult heart (Fig. 4B, 4C), and it has previously been reported that mammals undergo a postnatal catecholamine surge that is critical for cardiac maturation (37), we assessed whether expression of clock-controlled genes, as well as Sik1Intron and Nppbintron expression (to measure new transcription), increases specifically in response to α-adrenergic stimulation. Accordingly, NRVM were plated in culture medium containing 10% FCS for 24 hours, followed by serum starvation with medium supplemented with ITS for 48Hr, and subsequent treatment with the α-adrenergic receptor agonist PE (22) in serum-free medium for 4, 8, 12, 16, 20 and 24hrs (Fig. 5D). Consistent with the time scale on which Bmal1 has been shown to reach peak expression after norepinephrine treatment (11), Bmal1 transcript levels as measured by RT-qPCR peak after 8 hours of PE treatment, followed by gradual reversion of Bmal1 expression (Fig. 5E, left). Expression of Per2, an established Bmal1 target, in control-treated NRVM showed a significant peak at approximately 12-16Hr post treatment, relative to the trough of expression at 4-8 hours post treatment. In contrast, Per2 expression peaks at 8 hours post PE treatment, with a trough of expression appearing at 20 hours post PE treatment (Fig. 5E, right). These results support the notion that α-adrenergic stimuli can drive clock-controlled gene expression and alter the phase of their expression (11) (Fig. 5E, right, for the case of Per2). Along this same time course, PE induced robust hypertrophy and de novo transcription of Nppbintron (Fig. 5F). Under control conditions, a trough of Nppbintron was observed at 12 hours, and PE treatment resulted in overall increased levels of Nppbintron, with oscillation occurring within a 24-hour period (Fig. 5F, right). We observed that, similar to Bmal1, Sik1 reaches peak expression 4-8 hours of PE treatment, and this was followed by a gradual reversion of expression (Fig. 5G). Additionally, knockdown of Bmal1 decreased Sik1Intron transcript levels across at 0 and 2% FCS, with no significant difference at 10% FCS (Fig. 5H). Depletion of Bmal1 also impaired PE-mediated hypertrophy as indicated by cell size measurement and RT-qPCR for NppbIntron and impaired induction of Sik1Intron in response to PE (Fig. 5I&J).

Sik1 has been reported to be required for myocyte hypertrophy in response to a growth stimulus in mice (32). Intriguingly, we observed in rat neonatal myocytes that Sik1 knockdown alone significantly decreased myocyte size and total protein levels, accompanied by decreased NppbIntron and increased Per2Intron transcription. (Fig. 5K), highlighting roles for Sik1 in myocyte hypertrophy and tempering of clock-controlled gene transcription.

In summary, our results suggest that histone turnover is critical to myocyte hypertrophy, and that depletion of Bmal1 disrupts histone stoichiometry and impairs histone turnover, resulting in chromatin disorganization and increased accessibility of the Sik1 and Per2 promoters. In turn, this impairs myocyte hypertrophy in response to α-adrenergic stimulation (Fig. 5L).

## Discussion

In the study, we sought to determine whether Bmal1 acts to regulate histone turnover in the setting of neonatal cardiomyocyte growth. Our findings demonstrate that Bmal1 is required for cardiomyocyte hypertrophy in response to serum or α-adrenergic stimulation and that this effect involves chromatin remodeling at the Sik1 locus. A requirement for replication-independent turnover for myocyte hypertrophy is supported by our observation that knockdown of H3.3a, a replication-independent H3 variant (4,5), results in smaller myocytes and decreased Nppbintron transcription. Additionally, we characterized Sik1 as a novel clock-controlled gene, exemplified by its oscillatory expression in response to PE treatment, and demonstrated that Sik1 is required for hypertrophic gene transcription, as Sik1 knockdown impairs Nppbintron transcription and decreases myocyte size and total protein content.

Prior to this study, two seemingly contradictory models of Bmal1 knockout have been reported. One study reports early-onset dilated cardiomyopathy characterized by small cardiomyocytes in full-body Bmal1 knockout mice (38), while another study reports larger cardiac myocytes when Bmal1 is knocked out specifically in cardiac myocytes at E18.5 (18). Our data may reconcile these observations: we show that Bmal1 knockdown in the absence of external hypertrophic stimuli results in smaller cardiomyocytes and impaired Nppbintron transcription, whereas Bmal1 knockdown in myocytes in the presence of external hypertrophic stimuli (2-10% FCS) results in unchecked Nppbintron transcriptional activation. The results reported here, along with the previous reports described above, suggest that both timely external hypertrophic cues and a functioning myocyte clock are required for myocyte hypertrophy, and that the clock also regulates myocyte growth. As for how Bmal1 may both drive and regulate myocyte growth, previous studies have demonstrated that Bmal1 recruitment of cryptochrome proteins, Cry1 and Cry2 results in repression of Bmal1 targets (39). This observation indicates a potential mechanism awaiting further study in which Bmal1 and cryptochrome proteins antagonize transcriptional induction of Sik1, perhaps via antagonization of Gata4.

It has been previously reported that Bmal1 and Clock act as pioneer transcription factors (21), disrupting DNA-histone contacts to drive nucleosome dynamics and gene transcription. In the present study, Bmal1 knockdown prevented MNase digestion of Per2 and Sik1 promoter DNA, indicative of higher nucleosome occupancy or increased stability at these sites, and resulted in accumulation of H3 and H2A, of which H3.3 and H2A.Z are replication-independent histone variants that previous investigations have indicated are involved in hypertrophic gene transcription in neonatal myocytes (40). Accumulation of H3 and H2A, despite decreased translation indicated by decreased total cellular protein and decreased metabolic labeling of proteins found in the acid soluble chromatin fraction with Bmal1 knockdown, suggests a mechanism by which Bmal1 orchestrates ejection of histones targeted for degradation, as well as chromatinization of new histones. Further experimentation employing labeling of histones, followed by capture and sequencing of nucleosomes containing new histones (28), will yield spatiotemporal resolution on which loci, protein-coding or otherwise, are undergoing active turnover to allow for myocyte hypertrophy. Measurement of histone turnover across the genome, in conjunction with ChIP-Seq for various established histone post-translational modifications, will also increase our understanding of the epigenetic code behind what dictates histone turnover, and whether turnover occurs at transcriptionally silent as well as active chromatin (3,20).

When performing the gene ontology analysis to identify genes regulated by Bmal1 in a cardiac-specific manner, only Sik1 displayed two Bmal1 binding sites in its promoter, suggesting this regulatory axis is critical to maintenance of cardiac function. Recently, Sik1 has been shown to control the rate of clock re-entrainment in response to experimental jetlag, by tempering induction of clock-controlled genes. Specifically, behavioral responses were measured via actigraphy of control and Sik1-silenced mice subjected to an experimental jet-lag protocol of re-entrainment to a 6-hour phase advanced light/dark cycle. Two days after siRNA administration the light/dark cycle was advanced 6 hours, and 10 days later, the cycle was advanced 6 hours again. Actograms showed that Sik1-silenced mice showed significantly faster re-entrainment of behavior when compared to the controls in both cycle disruptions. This faster re-entrainment was reflected in MEFs expressing per1::luc subjected to serum shock followed by pharmacological inhibitor of Sik1, where inhibition of Sik1 resulted in a phase delay, i.e., a delay in the peak of, bioluminescence driven by the per1 promoter. This phase delay is thought to prevent singularity behavior of the clock, which is characterized as phaseless and unpredictable clock-controlled gene expression (41). This is perhaps reflected in our data showing increased Per2 expression following Sik1 knockdown (Fig. 5K). In the model organisms *Drosophila pseudoobscura* and *Neurospora crassa*, singularity behavior has been characterized as unpredictable or inappropriate initiation of development and cell proliferation in response to photic stimuli, respectively(41,42). We observe that in response to chronic pressure overload, Sik1 accessibility and expression decreases, suggesting that during cardiac pathology Bmal1-mediated chromatin organization of Sik1 is disrupted. This may translate to ill-timed modulation of clock-controlled growth, metabolism, or contractility in response to various external time cues described earlier, and result in downstream cardiac pathologies.

In summary, we provide evidence that Bmal1 represents a nexus point of nucleosome proteostasis by coordinating histone turnover and mediating chromatin accessibility at its gene targets. We also demonstrate that Sik1 represents a novel clock-controlled gene that has been implicated in both myocyte hypertrophy and clock entrainment that possesses a unique Bmal1-dependent signature of chromatin organization. This work sets the stage for future studies interrogating how Bmal1 orchestrates nucleosome proteostasis in postnatal cardiac myocytes *in vivo*, which are capable of both replication-dependent and independent forms of histone turnover while maintaining their unique epigenetic memory.

## Author Contributions

A.A., T.M.V. conceived of research

K.S., T.M.V. provided expertise, funding and infrastructure

A.A., A.R., T.K., M.R.G. performed experiments

A.A., D.J.C., A.R., T.M.V. analyzed data and interpreted results

A.A., D.J.C. prepared figures

A.A., T.M.V. wrote manuscript

All authors approved final version of manuscript.

## Methods

### Isolation of neonatal rat ventricular cardiomyocytes (NRVM) from P0 rat hearts

Myocytes were isolated as follows. Four litters of neonatal pups (approximately 48 animals, P2-3) were euthanized by decapitation, and atria were removed from excised hearts. Ventricles were then briefly rinsed in ice-cold 1x ADS buffer (116mM NaCl, 18mM HEPES, 845µM NaHPO4, 5.55mM Glucose, 5.37mM KCl, 831µM MgSO4, 0.002% Phenol Red, pH 7.35±0.5) and minced via ∼200 cuts with sterile micro scissors. Minced ventricles were then subjected to 5-10 serial digestions in a Wheaton 356945 Celstir 50mL Jacketed Glass Spinner Flask with Double Sidearms at room temperature (21°C) using 300μL/heart of ADS buffer containing 0.1% Trypsin (SigmaAldrich, Cat. No. T4799) and 0.002% DNase (Worthington Biochemical Corp., Cat. No LS002006). Each digestion lasted 10-20 minutes. The cell suspension from the first digestion was discarded as it contained unwanted dead cells and red blood cells. The cell suspension from each subsequent digestion was then combined with FCS to a concentration of 50% FCS. The pooled cell suspensions were then filtered with a 100µm nylon cell strainer (Foxx Life Sciences, Cat. No. 410-0003-OEM) and then centrifuged for 8 minutes at 1500xg. The resulting cell pellet was then resuspended in 8mL of 1×ADS buffer. Stock Percoll was prepared by combining 9 parts of Percoll (cat# 17-0891-02, GE healthcare) with 1 part of clear (without phenol red) 10×ADS. The stock Percoll was used to make the Percoll for the top (density= 1.059 g/ml; 1 part Percoll stock added to 1.2 parts 1x ADS without phenol red) and bottom (density= 1.082 g/ml; 1 part Percoll stock added to 0.54 parts 1x ADS with phenol red) layers. The gradient, consisting of 4 ml top Percoll and 3 ml bottom Percoll, was set in a 15 ml conical tube by pipetting the top Percoll first, and layering the bottom Percoll gently underneath. The cells (in 2 ml red 1x ADS buffer) were layered on the top of the discontinuous percoll gradient (4 gradients in total) and centrifuged at 1500xg for 30 minutes at 4°C, with no deceleration break to separate the myocytes from non-myocytes. The myocytes, concentrated between the two percoll layers as well as myocytes that centrifuged to the bottom of the tube, were then collected, washed once with 1x ADS buffer, and resuspended in plating medium composed of DMEM/F:12 (ThermoFisher Scientific, Cat. No. 11330-32) supplemented with 10% FCS and penicillin/streptomycin.

### Plating of NRVM

Fibronectin (ThermoFisher Scientific, Cat No. 33016015) dissolved at 1mg/mL in DNase/RNase-free molecular grade H2O was diluted to 5ug/mL in DMEM/F:12 containing penicillin/streptomyocin, and then added at 1mL/well to 6-well dishes (Corning, Cat. No. 3516), which were then incubated at 37°C for 30 minutes. The fibronectin solution was aspirated from the dishes, and percoll-purified myocytes suspended at 175,000 cells/mL in plating medium were then cultured on the fibronectin-treated 6-well dishes at 350,000 cells/well, 2mL of media/well. Cultures were then incubated at 37°C in an incubator infused with 95% N2/5% CO2 for 24 hours. *siRNA Transfection of Neonatal Rat Ventricular Myocytes:* 20uM stock siRNAs (diluted in DNase/RNase-free molecular grade H2O) were diluted to a concentration of 120nM using antibiotic-free DMEM/F:12 supplemented with 0.5% v/v. 24 hours after plating, the plating medium was aspirated and replaced with 1mL of the resulting siRNA solutions, which were incubated for 15 minutes at room temperature prior to being added to the culture dishes. Cultures were then incubated at 37°C in an incubator infused with 95% N2/5% CO2 for 7Hr, followed by replacement of transfection media with DMEM/F:12 supplemented with penicillin/streptomycin and 0, 2, or 10% FCS where indicated. Cultures treated with 0% FCS were supplemented with ITS (Corning, Cat. No. 354350) to maintain cell viability in the absence of serum.

### Treatment of NRVM with phenylephrine

To stimulate growth via phenylephrine, NRVM were first serum-starved for 48Hr with serum-free DMEM/F:12 supplemented with penicillin/streptomycin and ITS, and then treated with 50uM phenylephrine (Sigma-Aldrich, Cat. No. P6126) dissolved in DMEM/F:12 supplemented with ITS and antibiotics for 4-24Hr.

### siRNA sequences were purchased from Integrated DNA Technologies (IDT)

Negative Control DsiRNA, 5 nanomoles, Cat. No. 51-01-14-04 Bmal1 siRNA 1: 5’-rArGrUrArGrArArUrArCrArUrUrGrUrCrUrCrArArCrCrAAC-3’ Bmal1 siRNA 2: 5’-rCrArUrCrCrArArArArGrArUrArUrUrGrCrCrArArArGrUTA-3’ Sik1 siRNA 1: 5’-rGrCrUrArUrUrArArGrGrUrArCrUrArGrArArUrUrGrArUAA-3’

### Whole Cell Lysis

Medium was removed from 6-well culture dishes and adherent cells were washed twice with 1mL/well ice-cold DPBS and then lysed with 40-70µL of whole cell lysis buffer composed of 50 mM Tris (pH 8), 10mM EDTA, 1% SDS, 1x protease/phosphatase inhibitor mixture (Roche Applied Science; Catalog No. 05892791001 and 4906837001). Whole cell lysates were scraped and transferred to microcentrifuge tubes and stored at -80°C. Lysates were subsequently thawed and briefly vortexed, followed by clarification of lysate via centrifugation at 20,000xg for 5 minutes. Protein concentration of clarified cell lysates was measured via BCA protein assay kit (ThermoScientific, Cat. No. J63283.QA) using BSA dissolved in H2O as a standard.

### Subnuclear fractionation

To isolate nuclei, NRVM cultures (6 wells per sample) were washed twice with 1mL/well ice-cold PBS and lysed with 100µL of lysis buffer composed of 10 mM Tris (pH 7.4), 250 mM sucrose, 1 mM EDTA, 0.15% Nonidet P-40subsitute, and protease/phosphatase inhibitors, and scraped into Eppendorf tubes kept on ice. Samples were then centrifuged at 1000xg for 10 minutes, yielding a crude nuclear pellet. The crude nuclear pellet was then resuspended in 100µL and layered on top of a 300µL sucrose cushion composed on 1.6M sucrose, 15mM NaCl, 10mM Tris pH 7.4, and protease/phosphatase inhibitors. This was followed by centrifugation at 7500xg for 10 minutes. The resulting nuclear pellet was then washed once with lysis buffer composed of 10 mM Tris (pH 7.4), 250 mM sucrose, 1 mM EDTA, 0.15% Nonidet P-40subsitute, and protease/phosphatase inhibitors and centrifuged at 7500xg for 5 minutes. Chromatin was isolated from nuclei by resuspending nuclei in lysis buffer composed of 20mM HEPES, 7.5mM MgCl2, 30mM NaCl, 1M Urea, 1% Nonidet P-40subsitute, and protease/phosphatase inhibitors. This was followed by centrifugation at 13,000xg for 15 minutes, resulting in a supernatant containing the nucleoplasm and solubilized nuclear membranes and a chromatin pellet. The chromatin pellet was then incubated at 37°C for 16Hr in 100uL of 0.4N H2SO4. Samples were then centrifuged at 15,000xg for 10 minutes and the supernatant was transferred to a new tube and mixed with 33uL of 6.1N Trichloroacetic acid and kept on ice for 30 minutes to precipitate the acid soluble chromatin fraction. This was followed by centrifugation at 20,000xg for 20 minutes at 4°C. The supernatant was discarded while the pellet was washed once with 250µL ice-cold acetone and allowed to air dry. Pellets were then dissolved in 10-20µL cell lysis buffer composed of 50 mM Tris (pH 8), 10mM EDTA, 1% SDS, 1x protease/phosphatase inhibitor and quantified via BCA assay as described above.

### Covalent attachment of tags to capture histones and identify turnover over (CATCH-IT)

Fibroblasts isolated from P3 rat hearts were plated at 200,000 cells/well on 6-well dishes in DMEM/F:12 supplemented with 10% FCS and antibiotics and incubated overnight (∼16Hr). Cultures were then switched to serum-free DMEM/F:12 supplemented with ITS and antibiotics for 72Hr to halt cell cycling, and then switched to DMEM/F:12 supplemented with 20% FCS and antibiotics to stimulate synchronous cell cycling (PMID: 8491378). Cultures were then treated with L-azidohomoalanine hydrochloride (AHA) (Sigma-Aldrich, Cat. No. 900892) at a concentration of 4mM at 5Hr intervals after switching media to 20%. Cultures were then subjected to acid extraction of histones as described above. 10-30µg of acid soluble chromatin fraction were then pre-cleared with 50µL of pre-equilibrated streptavidin M280 Dynabeads (ThermoFisher,Cat. No. 11205D), and then subjected to click chemistry by adding the following reagents to the stated final concentrations: 300µM THPTA (Sigma-Aldrich, Cat. No. 762342), 750µM Ascorbic acid (Sigma-Aldrich, Cat.No. AX1775), 400µM CuSO 4 (Sigma-Aldrich, Cat. No. C1297), 7.5µM Biotin-PEG4-alkyne (Sigma-Aldrich, Cat. No. 764213). The reactions were incubated at room temperature for 1.5Hr and then precipitated with TCA as described above to remove excess click chemistry reagents. Precipitated protein was resuspended in 1% SDS, 15mM NaCl, 10mM Tris pH 7.4, and incubated with 100µL of pre-equilibrated streptavidin M280 Dynabeads at room temperature for 1.5Hr with agitation every 15 minutes. The beads were washed 4 times with 500 µL with 1% SDS, 15mM NaCl, 10mM Tris pH 7.4. To elute bound biotinylated proteins, beads were incubated at 100°C for 5 minutes in 30µL 1x NuPAGE LDS sample buffer (ThermoFisher, Cat. No. NP0007) supplemented with 25mM Biotin (PMID: 28986262) and 1.25% v/v β-mercaptoethanol. Eluted proteins were then resolved via SDS-PAGE on 15% poly-acrylamide gels and immunoblotted for histone H3 as described below. NRVM transfected with control or Bmal1-targeted siRNA as described above were maintained in serum-free medium supplemented with ITS for 72 hours. Following medium aspiration, cultures were treated with serum-free medium supplemented with ITS and 4mM AHA (1mL/well) and incubated at 37°C for 4Hr. Following isolation of acid soluble chromatin fractions as described above click reactions with Biotin-PEG4-alkyne were performed as described above using 250-500ng of protein, followed by SDS-PAGE and blotting with Streptavidin-HRP (Jackson Immunoresearch, Cat. No. 016-030-084).

### SDS-PAGE

SDS-PAGE electrophoresis buffer was composed of 24.2g Tris base, 115.2g Glycine, 8g sodium dodecyl sulfate, filled to a final volume of 8L. Immunoblot transfer buffer was composed of 24.2g Tris base, 115.2g Glycine, 1.28L methanol, filled to a final volume of 8L. SDS-PAGE gels were cast using mini-PROTEAN® spacer plates with 1.5 mm integrated spacers (Bio-Rad Cat. No. 1653312), mini-PROTEAN® short plates (Bio-Rad, Cat. No. 1653308), and 15-well 1.5mm mini-PROTEAN combs (Bio-Rad, Cat. No. 1653366).

### Total Protein Staining

Where indicated, following SDS-PAGE of 500-1000ng of acid soluble chromatin fractions isolated from NRVM, gels were stained with Oriole UV-fluorescent stain per the manufacturer’s instructions (Bio-Rad Cat. No. #1610496) to visualize and quantify total protein loading.

### Bmal1, GAPDH, and Cyclin A2 Immunoblots

5-7µg of whole cell protein lysates were subjected to SDS-PAGE at 200V for approximately 1 hour using 15% polyacrylamide gels. This was followed by electroelution onto PVDF membranes at 100V for 50 minutes under semi-dry transfer conditions at 4°C.

### H3, H4, H2B, H2A immunoblots

1µg of whole cell protein lysates or 100ng of acid soluble chromatin fractions were subjected to SDS-PAGE at 200V for 50 minutes on 15% polyacrylamide gels. This was followed by electroelution onto PVDF membranes at 100V for 50 minutes under semi-dry transfer conditions at 4°C.

Following electroelution, membranes were placed in methanol for 30 seconds and placed on paper towels to allow methanol to evaporate from the membranes. Membranes were then again placed in methanol for 30 seconds with gentle rocking, followed by placement into molecular grade H2O and then gently rocked for 12-16Hr 4 °C in phosphate-buffered saline supplemented with 0.01% Tween-20 (Sigma-Aldrich, Cat No. A7030) (PBST), 5% BSA (Cat. No. P2287), and the appropriate antibody. Membranes were then washed three times for 15 min in PBST and then incubated for 45 minutes at room temperature with the appropriate horseradish peroxidase– conjugated anti-IgG (Jackson ImmunoResearch Laboratories, Inc.) diluted at 1:2000 in 5% BSA dissolved in PBST. Membranes were then washed three times for 15 min with gentle rocking in TBST, subjected to enhanced chemiluminescence imaged using a BioRad ChemiDoc System. Immunoblots were quantified using ImageJ software densitometry.

### List of antibodies

| Target | Vendor | Cat. No. |
| --- | --- | --- |
| Bmal1 | Abcam | ab93806 |
| Gapdh | EMD Millipore-Sigma | MAB374 |
| H3 | Cell Signaling | 9715 |
| H2B | Abcam | ab1790 |
| H4 | Abcam | ab10158 |
| Cyclin A2 | Abcam | ab181591 |

### mRNA isolation

Medium was removed from 6-well culture dishes and adherent cells were washed with ice-cold DPBS and then lysed with 300µL of TRIzol™ Reagent (ThermoFisher Scientific, Cat. No. 15596026). The resulting lysates were homogenized and scraped into Eppendorf tubes and stored at -80°C. After thawing samples at room temperature (∼20°C), 60µL of chloroform were added to each sample, and shaken vigorously by hand for 15 seconds. Samples were then incubated at room temperature for 2-3 minutes to allow separation of layers. The samples were then centrifuged at 13,000rpm for 15 minutes at 4°C. The resulting upper clear layer (approximately 150uL) from each sample was pipetted into a fresh Eppendorf tube, and 150µL of isopropanol were added to each sample, followed by mixing via 10 inversions. The samples were then centrifuged at 13,000rpm for 20 minutes at 4°C. At this point, the samples are kept on ice, and the supernatant was removed without disturbing the visible pellet. 1mL of 75% reagent grade ethanol was then added to each tube, followed by centrifugation at 15,000rpm for 5 minutes at 4°C. The supernatant was discarded, followed by centrifugation at 15,000rpm for 5 minutes at 4°C, and any residual supernatant was also removed and discarded. The Eppendorf tubes were then opened and placed on a heat block set to 55°C for 5-10 minutes (or until no liquid remains). 10uL of DNase/RNase-free water was then added to the bottom of each tube, and the tubes were then closed and placed back on the heating block for 5 minutes to dissolve the RNA. This was then followed by centrifugation at 15,000rpm for 5 minutes at 4°C. The RNA concentration of the samples was then quantified by nanodrop.

### cDNA synthesis

Following quantification of mRNA concentration, 30-500ng of mRNA per sample was used to generate cDNA using an iScript™ cDNA Synthesis Kit (20µL reaction volume) using the following reaction protocol:

Priming: 5 min at 25°C

Reverse transcription: 50 min at 46°C

RT inactivation: 1 min at 95°C

Optional step: Hold at 4°C

Following cDNA synthesis, the cDNA synthesis reactions were diluted with RNase/DNase free H2O e.g., For every 500ng of mRNA used, 80µL of water was added to the cDNA reaction.

### RT-qPCR

Following dilution of cDNA reactions with water, target cDNA amplification was measured using the following reaction components: 5µL diluted cDNA, 1 µL of a 5 µM of a forward primer stock, 1 µL of a 5 µM reverse primer stock, 3 µL of RNase/DNase free H2O, and 10 µL SsoFast Evergreen Supermix (BioRad, Cat. No. 1725201).

### List of Reverse Transcription RT-qPCR primers

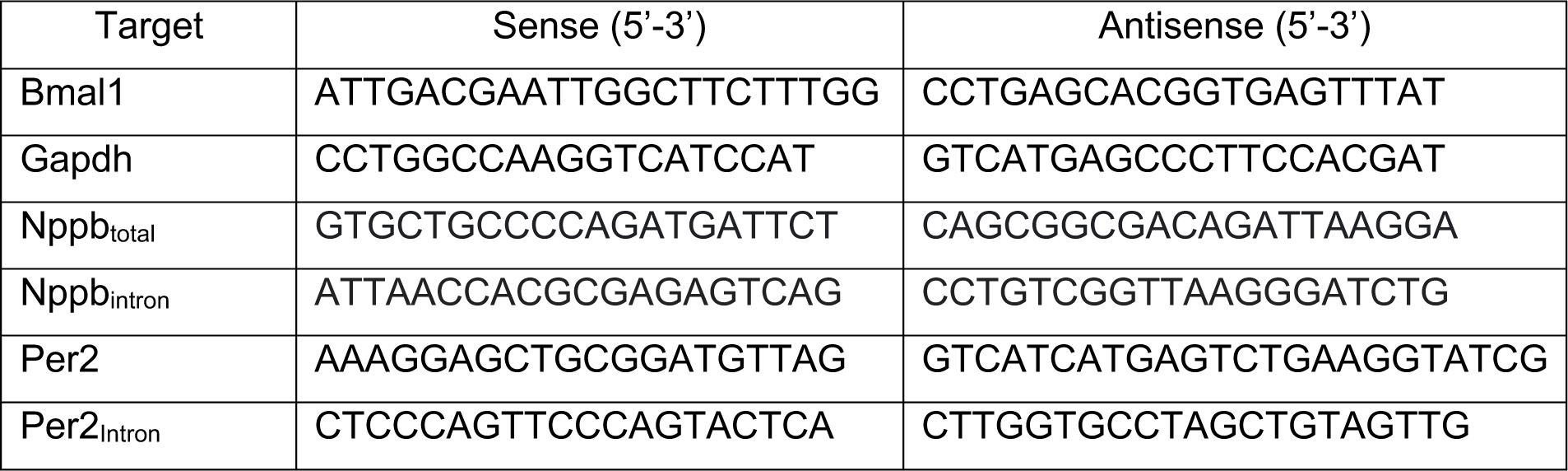
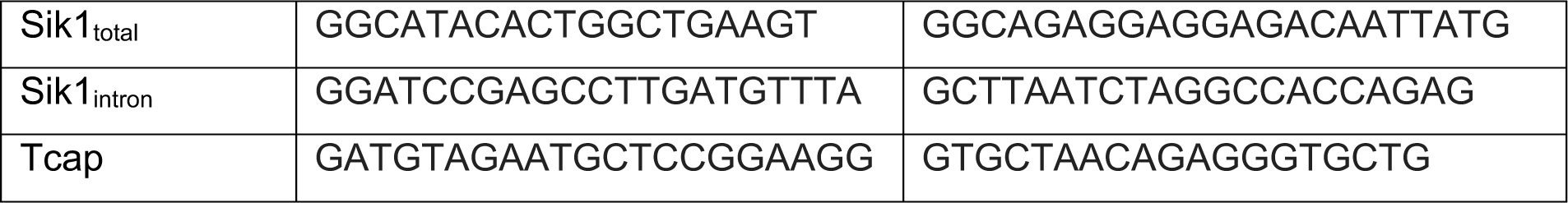

### MNase-qPCR(35)

Medium was removed from 6-well culture dishes and adherent cells were washed with ice-cold DPBS and then lysed with 100µL of ice-cold lysis buffer composed of 10 mM Tris-HCl, pH 7.4, 10 mM NaCl, 2 mM MgCl2, 0.5% Nonidet P-40Substitute and 1x protease/phosphatase inhibitor mixture. Lysates/intact nuclei were scraped and transferred to microcentrifuge tubes. For this specific protocol, each sample was composed of 2 wells scraped into the same tube. The resulting crude nuclear suspensions were then centrifuged at 1000xg for 10 minutes. At this point, the samples were maintained on ice. The supernatant was discarded, and the crude nuclear pellet was then gently but thoroughly resuspended in 120µL of pre-warmed MNase reaction buffer, followed by addition of 120µL of pre-warmed MNase reaction buffer (composed of 10mM Tris, pH 7.4, 5mM NaCl, 1mM CaCl2•2H2O) containing 0U, 0.001U, 0.01U, or 0.1U MNase (MNase, Worthington, LS004798). The reaction vessels were then immediately placed in a 37°C water bath for 5 minutes. This was followed by addition of 99µL of MNase stop reaction buffer composed of 60uL 100mM EDTA, 10mM EGTA, 30µL 20% SDS, and 9µL of 25mg/mL Proteinase K. The resulting solutions were then incubated for 16Hr at 37 °C. 340µL of phenol:chloroform:isoamyl alcohol (25:24:1) were then added to each and gently shaken by hand for 20 seconds. Samples were then centrifuged at 16,000xg at room temperature for 5 minutes. The upper aqueous layer was then transferred to a 2mL Eppendorf tube. To each the following were added in the stated order: 4.25µL of 15mg/mL Glyco-blue (ThermoFisher Scientific, Cat. No. AM9515), 170uL 7.5M ammonium acetate, and 1.27mL 100% reagent grade ethanol. Samples were then stored at -80°C for 1Hr. The samples were then centrifuged at 16,000xg for 30 minutes at 4°C to pellet DNA. The supernatant was discarded, and 150µL of 70% reagent grade ethanol was added, followed by centrifugation at 16,000xg for 2 minutes at 4°C. The supernatant was discarded, and another 150µL of 70% reagent grade ethanol was added, followed by centrifugation at 16,000xg for 2 minutes at 4°C. The supernatant was discarded, and the DNA pellets were allowed to air dry, followed by resuspension in 20µL DNase/RNase-free water and quantification by nanodrop. Chromatin accessibility was then assessed via RT-qPCR using the following reaction components: 5µL diluted Diluted gDNA (60ng/reaction), 1 µL of a 5 µM of a forward primer stock, 1 µL of a 5 µM reverse primer stock, 3 µL of RNase/DNase free H2O, and 10 µL SsoFast Evergreen Supermix.

### MNase-qPCR primers

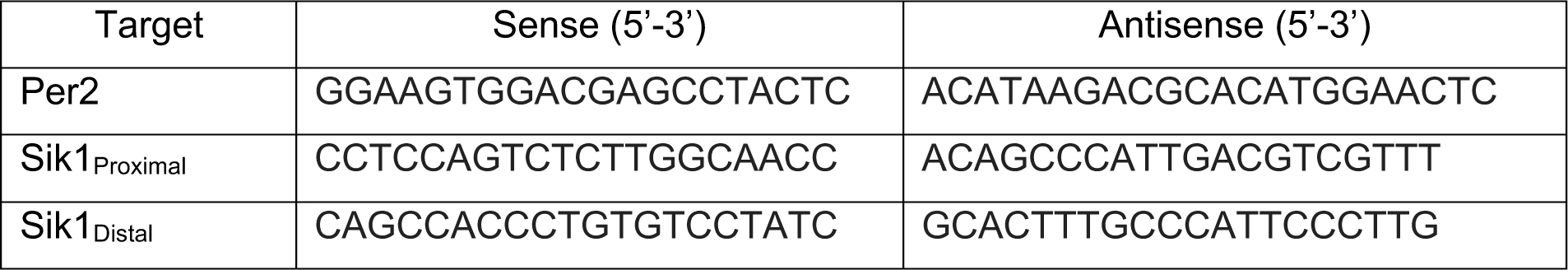

### Cell size measurements

NRVM cell size was measured via ImageJ. To accurately measure cell size in µm^2^, an image of a hemocytometer (FisherScientific, Cat. No. 02-671-6) was used to set the scale in ImageJ, and individual cells were manually traced on a Lenovo T460 Thinkpad with touchscreen.

### G:profiler analyses (43)

RNA-Seq data generated from adult and embryonic mouse myocytes in a previous study was downloaded into a new excel file and filtered for all genes significantly upregulated in adult myocytes (p<0.05). The resulting gene list was then filtered for genes containing Bmal1 ChIP-Seq peaks in their promoters/TSS found across organs, or for genes containing Bmal1 ChIP-Seq peaks in their promoters/TSS specific to the heart. The resulting gene lists were then subjected to gene ontology analysis via G:profiler analysis using default parameters.

### IGV Browser Track Viewing

The BW files(6,44) or bedgraph files(34) shown in Figure 5 were downloaded from NCBI gene expression omnibus and displayed on Integrated Genomics Viewer (IGV).

