## Supplemental Figure 1 for "Circadian Control of Histone Turnover During Cardiac Development and Growth"

### Figure S1

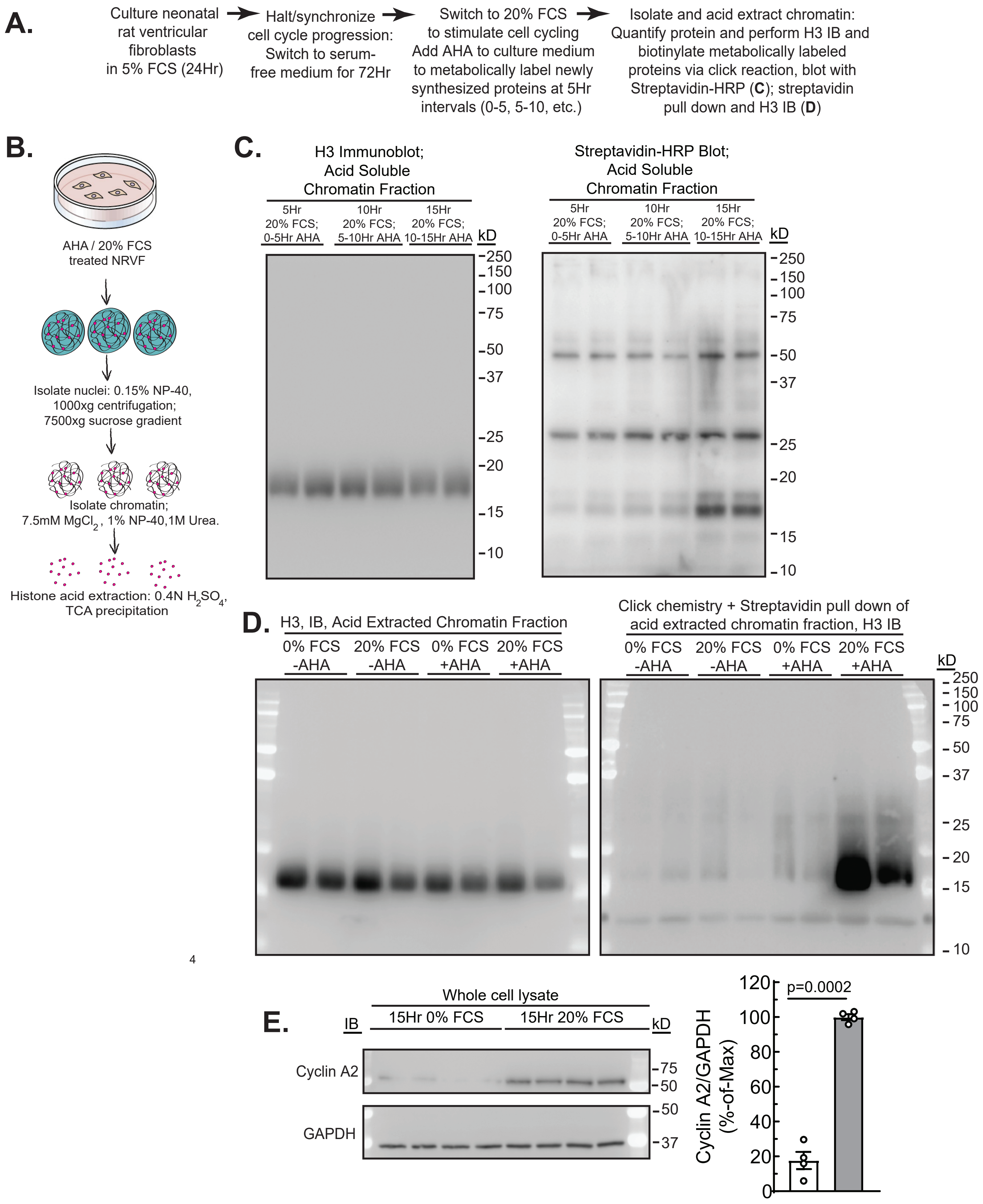
A.
Culture neonatal rat ventricular fibroblasts in 5% FCS (24Hr)
Halt/synchronize cell cycle progression: Switch to serum-free medium for 72Hr
Switch to 20% FCS to stimulate cell cycling Add AHA to culture medium to metabolically label newly synthesized proteins at 5Hr intervals (0-5, 5-10, etc.)
Isolate and acid extract chromatin: Quantify protein and perform H3 IB and biotinylate metabolically labeled proteins via click reaction, blot with Streptavidin-HRP (C); streptavidin pull down and H3 IB (D)
B.
AHA / 20% FCS treated NRVF
Isolate nuclei: 0.15% NP-40, 1000xg centrifugation; 7500xg sucrose gradient
Isolate chromatin; 7.5mM $MgCl_2$, 1% NP-40,1M Urea.
Histone acid extraction: 0.4N $H_2SO_4$, TCA precipitation
C.
H3 Immunoblot; Acid Soluble Chromatin Fraction
5Hr 20% FCS; 0-5Hr AHA
10Hr 20% FCS; 5-10Hr AHA
15Hr 20% FCS; 10-15Hr AHA
kD
250
150
100
75
50
37
25
20
15
10
Streptavidin-HRP Blot; Acid Soluble Chromatin Fraction
5Hr 20% FCS; 0-5Hr AHA
10Hr 20% FCS; 5-10Hr AHA
15Hr 20% FCS; 10-15Hr AHA
kD
250
150
100
75
50
37
25
20
15
10
D.
H3, IB, Acid Extracted Chromatin Fraction
0% FCS -AHA
20% FCS -AHA
0% FCS +AHA
20% FCS +AHA
Click chemistry + Streptavidin pull down of acid extracted chromatin fraction, H3 IB
0% FCS -AHA
20% FCS -AHA
0% FCS +AHA
20% FCS +AHA
kD
250
150
100
75
50
37
25
20
15
10
E.
IB
Whole cell lysate
15Hr 0% FCS
15Hr 20% FCS
kD
Cyclin A2
75
50
GAPDH
50
37
Cyclin A2/GAPDH (%-of-Max)
p=0.0002
0
20
40
60
80
100
120

**Supplemental Figure 1**: **AHA incorporation into total histone H3 in cell cycle synchronized neonatal rat ventricular fibroblasts (NRVF). A**) Experimental timeline of cultured NRVF subjected to cell cycle synchronization in combination with AHA labeling at different intervals after serum treatment. This was followed by acid extraction of chromatin-associated histones and subsequent detection of AHA-labeled proteins with alkyne-biotin and Streptavidin-HRP. **B**) Diagram illustrating the process of histone acid extraction from chromatin. **C**) H3 immunoblot (left) and Streptavidin-HRP blot of acid soluble chromatin fractions from NRVF treated with serum and 4mM AHA for the indicated times. **D**) H3 immunoblots of acid extracted chromatin fractions and concomitant streptavidin pulldowns of AHA-labeled proteins from NRVF treated with or without 20% FCS for 15Hr and with or without 4mM AHA for the last 5Hr. **E**) Cyclin A2 and GAPDH immunoblots indicating increased AHA labeling observed in **C** and **D** occurs during S-phase of the cell cycle. P-values indicated are generated by unpaired t-tests.
